# Imaging-based evaluation of pathogenicity by novel *DNM2 variants* associated with centronuclear myopathy

**DOI:** 10.1101/2021.03.01.433478

**Authors:** Kenshiro Fujise, Mariko Okubo, Tadashi Abe, Hiroshi Yamada, Kohji Takei, Ichizo Nishino, Tetsuya Takeda, Satoru Noguchi

## Abstract

Centronuclear myopathy (CNM) is characterized clinically by muscle weakness and pathologically by the presence of centralized nuclei and disarrangement of T-tubules in muscle fibers. *DNM2* which encodes a large GTPase dynamin 2 have been identified as a causative gene for CNM. Nevertheless, the identification of *DNM2* variants may not always lead to the definitive diagnosis as their pathogenicity is often unknown.

In this study, by imaging T-tubule-like structures reconstituted *in cellulo*, we demonstrated that aberrant membrane remodeling by mutant dynamin 2 is tightly associated with gain-of-function features of *DNM2* variants. This simple *in cellulo* assay provided quantitative data required for accurately evaluating pathogenicity of reported and novel *DNM2* variants identified from CNM patients in our cohort. Our approaches combining the *in cellulo* assay with clinical information of the patients enabled to explain the course of a disease progression by pathogenesis of each variant in *DNM2*-associated CNM.

## INTRODUCTION

Centronuclear myopathy (CNM) is a congenital myopathy characterized clinically by slowly progressive muscle weakness and pathologically by the presence of centralized myonuclei, radial arrangement of sarcoplasmic strands of oxidative enzymes, and type 1 fiber predominance and hypotrophy(Jungbluth, Wallgren-Pettersson, & Laporte, 2008). In addition, T-tubules (transverse tubules) and triads are disorganized on electron microscopic examination(Al-Qusairi & Laporte, 2011). So far, seven genes, *MTM1*, *SPEG*, *BIN1, DNM*2, *RYR1*, *TTN* and *CCDC78*, have been reported to be causative for CNM(Agrawal et al., 2014; Jungbluth & Gautel, 2014; Majczenko et al., 2012; Romero, 2010). Diverse clinical manifestations among CNM patients are mainly attributed to the different responsible genes and mutations. In *DNM2*, 26 pathogenic variants have been reported to cause the autosomal dominant form of CNM with relatively mild and slowly progressive symptoms(Biancalana et al., 2018; Bohm et al., 2012; Casar-Borota et al., 2015; Hohendahl, Roux, & Galli, 2016). *DNM2* encodes the ubiquitous isoform of dynamin, dynamin 2, which is a GTPase essential for membrane fission in endocytosis(Antonny et al., 2016; Ferguson & De Camilli, 2012). The defects in either self-assembly or membrane binding ability of dynamin 2 should be the common cause of *DNM2*-associated CNM since most of the reported pathogenic *DNM2* variants were missense in the stalk and pleckstrin homology (PH) domain.

A huge number of variants are now identified for various congenital diseases by massive parallel sequencing technologies but pathogenicity is not unknown for many of them mostly due to lack of functional testing. This is also the case with *DNM2*-associated CNM as functional assays, such as GTPase measurement, self- assembly and membrane binding assays of the mutated dynamin, which may well determine pathogenicity of the identified variants, are not always accessible without special settings.

In this study, pathogenicity of both reported and novel *DNM2* variants associated with CNM were systematically and quantitatively analysed by imaging T-tubule like structures (TLS) reconstituted *in cellulo*. Our approaches combining the *in cellulo* assay with clinical information clearly demonstrated strong correlation between genotypes, cellulotypes and disease phenotypes in *DNM2*-associated CNM. These results suggest that aberrant membrane remodeling by *DNM2* variants is tightly linked to the pathogenesis and prognosis of CNM.

## MATERIALS AND METHODS

### Editorial Policies and Ethical Considerations

National Center of Neurology and Psychiatry (NCNP) has been functioning as a referral center for muscle disease since 1978. All the samples and clinical data used in this study were sent to NCNP from the physicians for diagnostic purposes (until March 2019). Written consent was obtained from parents or guardians. This study was approved by the Ethics Committee in NCNP.

### Genetical and histological analyses of patients

Genetic variants were analysed using genomic DNA from serum or biopsied muscles from 3933 cases which was suspected to be muscle diseases. Genetic analyses were performed by targeted re-sequencing that covered all exonic regions and exon-intron borders in *DNM2* gene (2858 analyses) and/or by whole exome sequencing (2599 analyses) as described previously (Nishikawa, Mitsuhashi, Miyata, & Nishino, 2017). Histological analysis was performed using muscle samples taken from the biceps brachii and then frozen in isopentane cooled in liquid nitrogen as described previously (Okubo et al., 2018).

### Plasmid construction, cell culture and DNA transfection

All the expression constructs used in this study were generated using Gateway Cloning Technology (Thermo Fisher Scientific). Entry clones of human dynamin 2 (NM_001005360) and human BIN1 isoform 8 (NM_004305.4) were prepared by B-P recombination cloning of PCR products respectively amplified from pcDNA3.1-GFP-Topo-hDNM2-WT (generous gift from P. Guicheney, UPMC) and pEGFP-mAmph2 (generous gift from P. De Camili, Yale University) using corresponding primers (BIN1 fw: 5’-ggggacaagtttgtacaaaaaagcaggctgcatggcagagatgggcag-3’; BIN1 rv: 5’-ggggaccactttgtacaagaaagctgggtctgggaccctctcagtgaag-3’, Dynamin 2 fw: 5’-ggggacaagtttgtacaaaaaagcaggctgcatgggcaaccgcggga-3’, Dynamin 2 rv: 5’-ggggaccactttgtacaagaaagctgggtcgtcgagcagggatggc-3’) into pDONR201 vector. Expression constructs of dynamin 2 and BIN1 were prepared by L-R recombination cloning of their Entry clones into Destination vectors (generous gift from H. McMahon, MRC-LMB) either for expressing proteins in mammalian cells (pCI vectors for expressing FLAG-, or RFP-tagged proteins) or for bacterial protein expression (pET15b for His-fusions and pGEX-6P-2 for GST-fusions) (generous gift from H. McMahon, MRC-LMB).

C2C12 cells (ATCC CRL-1722) was grown in D-MEM (High Glucose) with L-Glutamine, Phenol Red and Sodium Pyruvate (FUJIFILM Wako chemicals, 043-30085) supplemented with 10% fetal bovine serum (FBS) (Gibco, 12483, Lot No.1010399) and Penicillin-Streptomycin (Gibco, 15140122) at 37 °C in 5% CO_2_. For transfection of C2C12, 70% confluent cells in VIOLAMO VTC-P24 24-well plates (AS ONE, 2-8588-03) were transfected with 0.5 *μ*g expression plasmids using Lipofectamine LTX with Plus Reagent (Thermo Fisher Scientific, 15338100). To examine consequences of the expression of BIN1 or dynamin 2 in either wild type (WT) or mutant forms, cells were fixed after 48 h of the transfection for phenotypic analyses.

### Introduction of CNM mutations into dynamin 2

Entry clones for the CNM mutants of dynamin-2 (R465W) was prepared by BP recombination reaction of PCR products amplified from pcDNA3.1-GFP-Topo-hDNM2-R465W (generous gift from P. Guicheney, UPMC) using corresponding primers (Supplementary Table 1) into pDONR201. Entry clones for other mutant dynamin 2 were prepared by introducing corresponding mutations into the Entry clone of wild type human dynamin 2 using QuikChange Lightning Site-directed Mutagenesis kit (Agilent Technologies, 210518) following manufacturer’s instruction. Sense and antisense primers used for the site-directed mutagenesis are as follows.

E368K sense: 5’-tcaatcgcatcttccacaagcggttcccatttgag-3’
E368K antisense: 5’-ctcaaatgggaaccgcttgtggaagatgcgattga-3’
R369Q sense: 5’-cgcatcttccacgagcagttcccatttgagctg-3’
R369Q antisense: 5’-cagctcaaatgggaactgctcgtggaagatgcg-3’
S619L sense: 5’-cagctggaaggccttgttcctccgagctg-3’
S619L antisense: 5’-cagctcggaggaacaaggccttccagctg-3’
G495R sense: 5’-ccatgaggacttcatcaggtttgccaatgccca-3’
G495R antisense: 5’-tgggcattggcaaacctgatgaagtcctcatgg-3’
V520G sense: 5’-gggagatcctggggatccgcagggg-3’
V520G antisense: 5’-cccctgcggatccccaggatctccc-3’
G624V sense: 5’-ttcctccgagctgtcgtctaccccgag-3’
G624V antisense: 5’-ctcggggtagacgacagctcggaggaa-3’
P294L sense: 5’-gggagtcgctgctggccctacgtag-3’
P294L antisense: 5’-ctacgtagggccagcagcgactccc-3’
R724H sense: 5’-ggacgacatgctgcacatgtaccatgccc-3’
R724H antisense: 5’-gggcatggtacatgtgcagcatgtcgtcc-3’

### Immunofluorescent microscopy and quantitative analysis of TLS

Primary antibodies used in this study were polyclonal rabbit anti-DDDDK tag (MBL, PM020). The secondly antibody used in this study, Alexa Fluor 488-conjugated donkey anti-Rabbit IgG (H+L) (A21206), was purchased from Thermo Fisher Scientific. For immunostaining of C2C12, cells grown on coverslips were fixed with 4% paraformaldehyde (EMS, 15710) in PBS for 15 min at room temperature. After washing with PBSTB (PBS containing 0.1% Triton X-100, 1% BSA), the cells were permeabilized and blocked with PBS containing 0.5% Triton X-100 and 3% BSA for 1 h at room temperature. The samples were then incubated with primary antibodies diluted 1:1000 in PBSTB overnight at 4 °C in a humid chamber. After washing with PBSTB, the cells were incubated with secondly antibodies diluted in PBSTB for 3 h at room temperature. Then, the cells were washed with PBSTB and mounted in Fluoromount/Plus (K048, Diagnostic BioSystems). Immunostained cells were observed under BX51 fluorescence microscope (OLYMPUS) and images were acquired with Discovery MH15 CMOS camera and ISCapture image acquisition software (Tucsen). All images were analyzed using FIJI (Schindelin et al., 2012) and processed with Photoshop (Adobe).

### Quantitative analysis of TLS

Quantitative analysis of the TLS was performed by FIJI as described previously (Fujise et al., 2020). Firstly, background signal was subtracted from microscopic images of BIN1-expressing cells (Rolling ball radius = 10 pixels). Then, the membrane tubules were enhanced with FFT Bandpass Filter (Filter: large structures down to 5 pixels and up to 3 pixels; Suppress stripes: None; Tolerance of direction: 5%). The membrane tubules were detected and binarized with Threshold command and the binarized membrane tubules were skeletonized to be analyzed with Analyze Skeleton (2D/3D) plugin. Membrane tubules with length between 0.5 and 2 *μ*m were considered to be as “short”.

### In vitro sedimentation assay using recombinant BIN1 and dynamin 2

Recombinant protein of BIN1 isoform 8 and dynamin 2 was expressed and purified as GST fusion and His-tagged proteins, respectively as described previously (Fujise et al., 2020).

*In vitro* sedimentation assay of dynamin 2 was performed as described previously (Fujise et al., 2020). In short, wild type or CNM mutant (E368K, R369Q, R465W and S619L) dynamin 2 were diluted to 1 *μ*M in reaction buffer (10 mM Hepes, 2 mM MgCl_2_, 100 mM NaCl, pH 7.5) and incubated for 5 min at 37 ◻. To induce disassembly, 1 mM GTP was added to the preassembled dynamin 2 and incubated for 5 min at 37 ◻. The samples were centrifuged at 230,000*g* for 10 min at 25 ◻ using CS100GXL ultracentrifuge and S120AT3 rotor (Eppendorf Himac Technologies) and resultant supernatant and pellet were analyzed by SDS-PAGE followed by Coomasie Brilliant Blue R-250 staining.

### Dynamin GTPase activity

GTPase activity of dynamin 2 was determined by monitoring release of free orthophosphate using malachite green assay as described previously (Fujise et al., 2020). The malachite green reagent was prepared by mixing solution A (17 mg of Malachite Green Carbinol base dye (229105, Merck) in 20 mL 1 N HCl) and Solution B (0.5 g Ammonium molybdate (277908, Merck) in 7 mL 4 N HCl) with filling up to 50 mL by MilliQ water followed by filtration through 0.45 *μ*m membrane (S-2504, KURABO). In the assay, 0.2◻ *μ*M dynamin in the presence of BIN1 at different molar ratio was mixed with 1 mM GTP in GTPase reaction buffer (10 mM Hepes, 2 mM MgCl_2_, 50 mM NaCl, pH 7.5) with or without 0.005 *μ*g/*μ*L lipid nanotubes and incubated for 5 min at 37 ◻. After the reaction was stopped on ice for 10 min, 160 *μ*L of malachite green reagent was added to the 40 *μ*L of the reaction mix in 96 well plate (442404, Thermo Fisher Scientific). After 5 min shaking at 1200 rpm with Digital MicroPlate Genie Pulse (Scientific Industries, Inc.), released orthophosphate was colorimetrically quantified by measuring OD 650 nm using a microplate reader (SH-1000, CORONA ELECTRIC).

### Statistical data analysis

Statistical data analysis was performed using Prism 8 (GraphPad Software) and Excel (Microsoft). For all quantification provided, means and SEM are shown. Statistical significance was determined using a two-sided *t* test and P values are shown in the figures.

### Data availability

All relevant data are included with the manuscript or available from the authors upon request.

## RESULTS

### Identification of SNVs from CNM patients by cohort analyses

We identified 17 sporadic patients with *DNM2* variants in 3933 cases who were suspected to have muscle diseases. Among these patients, 11 patients carried reported variants (Supplementary Table 1), while 6 had novel missense variants (Supplementary Table 2). In total, five novel heterozygous variants, c.1483G>A (p.G495R), c.1559T>G (p.V520G), c.1871G>T (p.G624V), c.881C>T (p.P294L) and c.2171G>A (p.R724H), were identified (Supplementary Table 2). The predicted substitutions in amino acid residues occurred either at the unstructured loops in PH domain (Val520 and Gly624) and stalk domains (Gly495) or at bundle signaling element domain, a flexible hinge between the G-domain and stalk (Pro294 and Arg724) based on the crystal structure of human dynamin 1 (Supplementary Fig. 1). Histological analyses of skeletal muscle biopsies from the patients with reported variants exhibited typical myotubular myopathic features (P2) or CNM pathological features (P1, P3-P11), including centrally placed nuclei, peripheral halo, presence of radial sarcoplasmic strands, type 1 fiber predominance and adipose tissue infiltration (Supplementary Fig. 2). Consistently, the typical clinicopathological features of CNM were observed in patients with the novel variants except for P16 and P17 (Supplementary Table 2).

### CNM variants induced gain-of-function features of dynamin 2 in vitro

To characterize pathogenicity of reported *DNM2* variants, we analyzed the mutant dynamins (E368K, R369Q, R465W and S619L) for their self-assembly and GTPase activities, both of which are essential for membrane fission by dynamin (Fujise et al., 2020; Marks et al., 2001; Ramachandran et al., 2007; Wang et al., 2010; Warnock, Hinshaw, & Schmid, 1996). In the sedimentation assay, purified wild type and mutant dynamin 2 self-assembled in the absence of GTP and more than 90% of proteins are recovered in the precipitate (Fig. 1A and Supplementary Fig. 3, -GTP). Previous studies demonstrated that CNM mutants of dynamin 2 self-assemble to form stable polymers resistant to GTP hydrolysis-dependent disassembly (Fujise et al., 2020; Ramachandran et al., 2007; Wang et al., 2010). Consistently, almost all the mutant dynamin 2 remained in the precipitate even after GTP addition (Fig. 1A and Supplementary Fig. 3, E368K, R369Q, R465W and S619L, +GTP). In contrast, more than 30% of self-assembled wild type dynamin 2 were disassembled and recovered in the supernatant after GTP addition (Fig. 1A and Supplementary Fig. 3, WT, +GTP).

**Figure 1.**
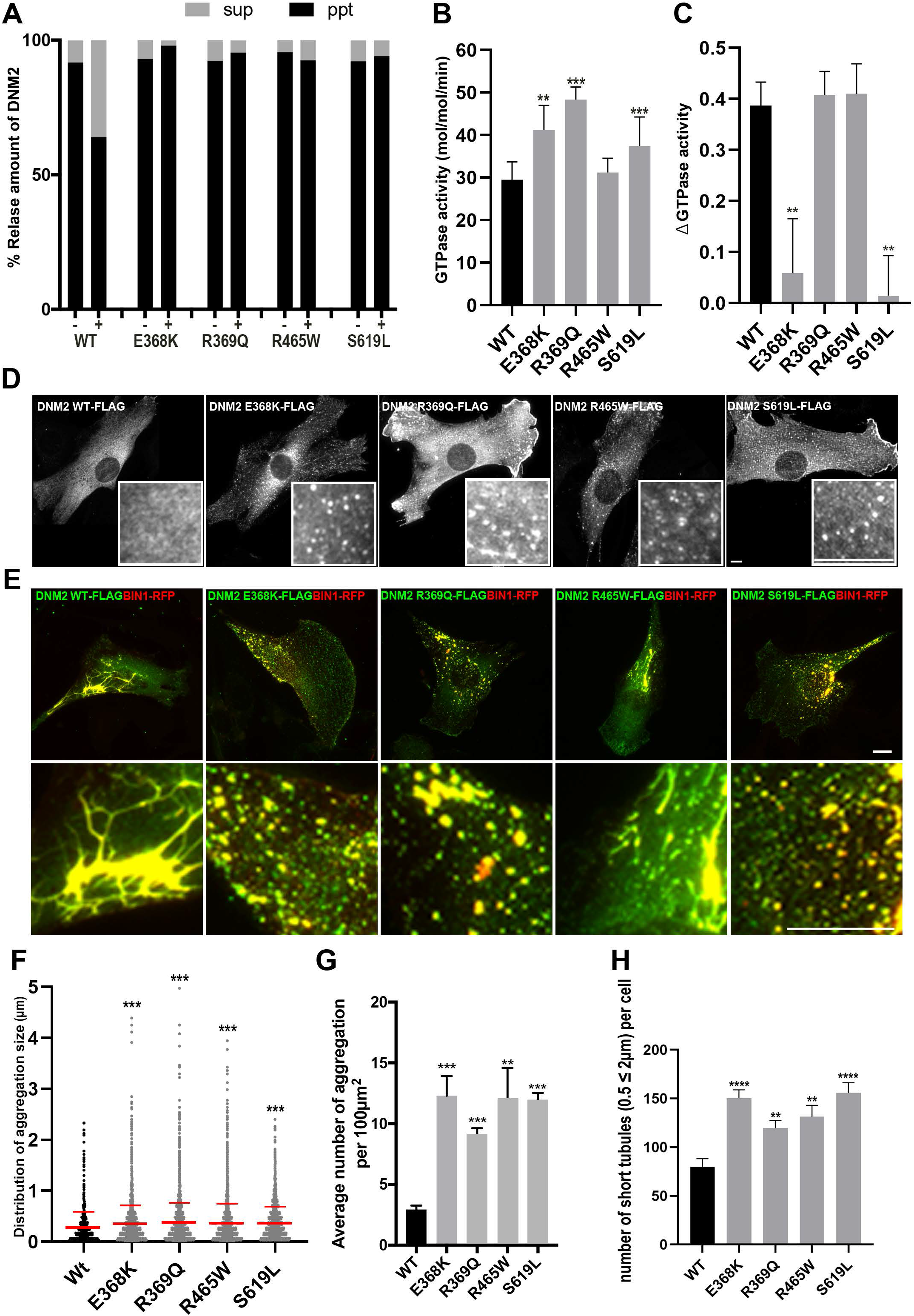
Gain of function features of mutant dynamin 2 in vitro and in cellulo. (A) Quantitative analysis of the *in vitro* sedimentation assay. Relative amount of wild type (WT) or mutant dynamin 2 (E368K, R369Q, R465W and S619L) either in the supernatant (sup) or in the precipitate (ppt) with or without GTP (+ or −) are shown. (B) GTPase activity of wild type (WT) and mutant dynamin 2 (E368K, R369Q, R465W and S619L). Data are means ± SEM (n=3, N=3). (C) BIN1-mediated inhibition of dynamin 2 GTPase activity. Relative ratios of inhibited GTPase activities (ΔGTPase activity) of wild type (WT) and mutant dynamin 2 (E368K, R369Q, R465W and S619L) in the presence of BIN1 (dynamin 2: BIN1 = 1:4 in molar ratio) are shown. Data are means ± SEM (n=3, N=3). (D) Formation of aggregates by mutant dynamin 2. Localization of FLAG-tagged wild type dynamin 2 (DNM2 WT-FLAG) or mutant dynamin 2 (DNM2 E368K-FLAG, DNM2 R369Q-FLAG, DNM2 R465W-FLAG and DNM2 S619L-FLAG) in C2C12 cells are shown. Scale bars are 10 *μ*m. (E) TLS formation in the presence of wild type and CNM mutant dynamin 2. Merged images of FLAG-tagged wild type (DNM2 WT-FLAG) or mutant dynamin 2 (DNM2 E368K-FLAG, DNM2 R369Q-FLAG, DNM2 R465W-FLAG and DNM2 S619L-FLAG) (green) with BIN1-RFP (red) are shown. Scale bars are 10 *μ*m. (F) Enhanced aggregate formation by mutant dynamin 2. Size of aggregates formed by either wild type or mutant dynamin 2 (shown in D) are measured and their distribution is shown. Data are means ± SEM (n ≥ 891 aggregates in 10 cells). (G) Increased number of aggregates by mutant dynamin 2. Average number of the aggregates formed either by wild type or by mutant dynamin 2 per 100 μm^2^ of cell area are shown. Data are means ± SEM (n ≥ 967 aggregates in ≥ 10 cells, N=3). (H) Quantification of the short TLS (0.5 ≤ 2 *μ*m). Data are means ± SEM (n ≥ 936 TLS in ≥ 10 cells, N=3).

Previous studies showed that elevated GTPase activity is a character of mutant dynamin 2 with CNM variants (Chin et al., 2015; Fujise et al., 2020; Kenniston & Lemmon, 2010; Wang et al., 2010). Consistent with the previous studies, E368K, R369Q, and S619L mutants exhibited higher GTPase activities compared to that of wild type dynamin 2 (Fig. 1B). However, statistical significance of the elevated GTPase activity was not confirmed for R465W mutant (Fig. 1B). Previous studies showed that BIN1 is a negative regulator of dynamin 2 (Cowling et al., 2017; Fujise et al., 2020). Consistently, GTPase activity of wild type dynamin 2 was stoichiometrically inhibited by BIN1 (Fig. 1C and Supplementary Fig. 4, WT). In contrast, BIN1 failed to inhibit GTPase activities of some mutant dynamin 2 (E368K and S619L), but, interestingly, those of other mutants (R369Q and R465W) were inhibited by BIN1 (Fig. 1C and Supplementary Fig. 4). These *in vitro* data suggest that CNM mutants generally exhibit elevated GTPase activity and they are resistant to BIN1-mediated inhibition, although existence of the exceptional variants in GTPase activation and BIN1 sensitivity suggest limitation of the experimental approaches.

### Imaging-based evaluation of functional defects caused by genetic variants in DNM2

In C2C12 cells, FLAG-tagged wild type dynamin 2 formed very fine puncta, while all the mutant dynamin 2 with reported variants either in the stalk (E368K, R369Q and R465W) or in the PH domain (S619L) formed abnormally large puncta (Fig. 1D, F and G). We previously demonstrated that co-expression of dynamin 2 and BIN1 in C2C12 cells induce TLS, membranous tubular structures mimicking T-tubules in skeletal muscles (Fujise et al., 2020). Wild type dynamin 2 was recruited to BIN1 to induce thicker and unevenly distributed TLS (Fig. 1E, DNM2 WT-FLAG). In contrast, all the mutant dynamin 2 induced shorter dot-like TLS despite they are still colocalized with BIN1 (Fig. 1E). Quantitative analyses showed that the number of shorter TLS (0.5-2 *μ*m) was increased in the presence of mutant dynamin 2 (Fig. 1H).

### Dynamin 2 with CNM-associated novel variants induced shorter TLS

We next explored pathogenicity of the novel *DNM2* variants that cause P294L, G495R, V520G, G624V and R724H substitutions by analyzing their effects on TLS formation. Similar to the reported CNM mutants, FLAG-tagged proteins of the novel *DNM2* variants, G495R, V520G and G624V formed aggregates in C2C12 cells (Fig. 2A, C and D). Furthermore, these three mutants also induced significantly shorter TLS compared to those with wild type dynamin 2 (Figs. 2B and E). Interestingly, two novel *DNM2* variants, P294L and R724H, formed neither aggregates (Fig. 2A, C and D) nor aberrantly shorter TLS (Fig. 2B and E). These findings are compatible with the histological and clinical features of the patients P16 and P17, harboring c.881C>T (P294L) and c.2171G>A (R724H) variants, in which atypical CNM histopathology with low number of centrally placed nuclei devoid of radial strands were observed (Supplementary Fig. 2).

**Figure 2.**
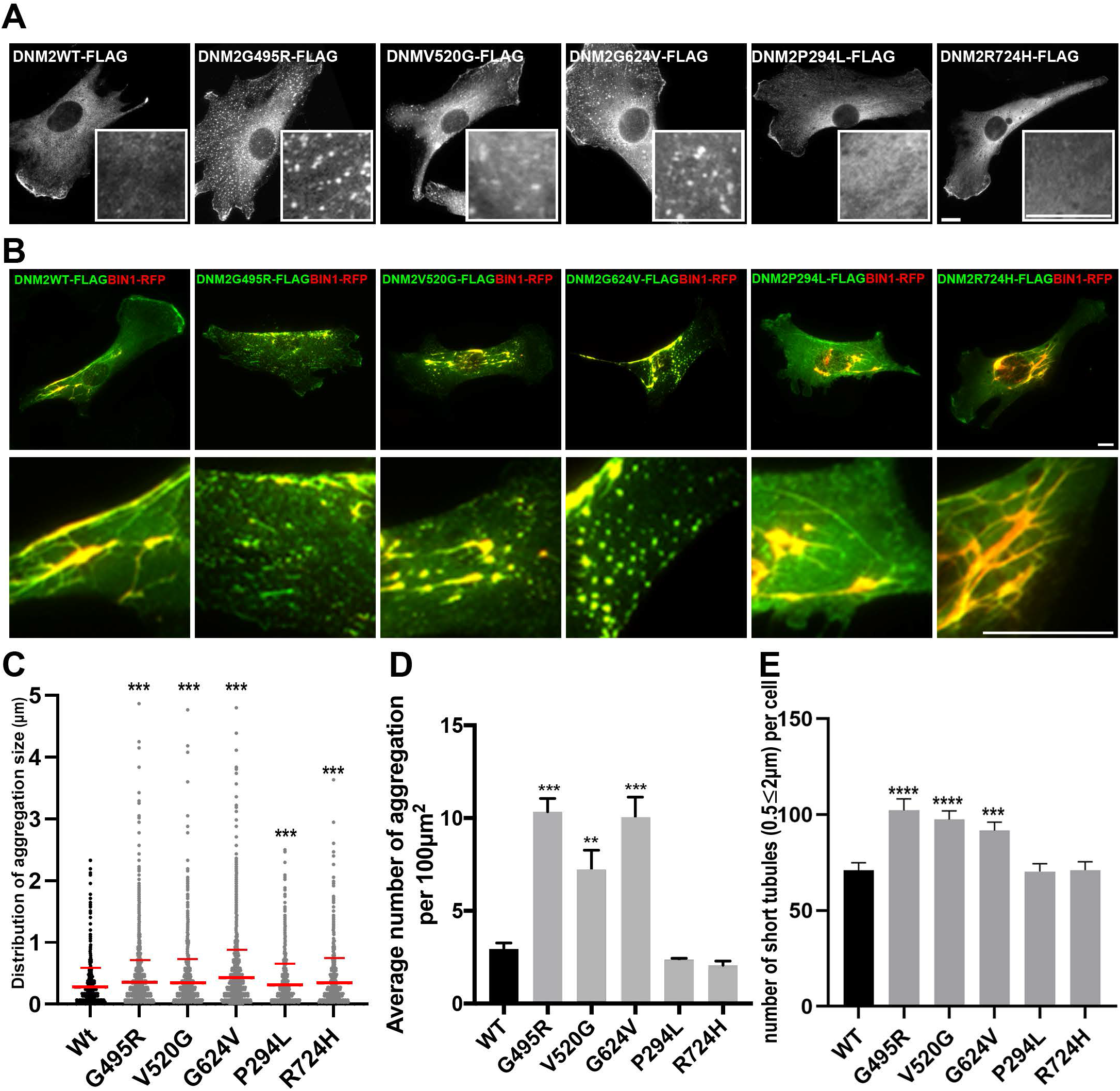
TLS formation by novel CNM-associated dynamin 2 mutants. (A) Aggregate formation by FLAG-tagged wild type dynamin 2 (DNM2 WT-FLAG) and five novel mutant dynamin 2 (DNM2 G495R-FLAG, DNM2 V520G-FLAG, DNM2 G624V-FLAG, DNM2 P294L-FLAG and DNM2 R724H-FLAG) in C2C12 cells. Scale bars are 10 *μ*m. (B) Defective TLS formation by novel mutant dynamin 2. Merged images of FLAG-tagged wild type (DNM2 WT-FLAG) or novel mutant dynamin 2 (DNM2 G495R-FLAG, DNM2 V520G-FLAG, DNM2 G624V-FLAG, DNM2 P294L-FLAG and DNM2 R724H-FLAG) (green) with BIN1-RFP (red) are shown. Scale bars are 10 *μ*m. (C) Distribution of the average diameter of aggregates formed by either wild type or mutant dynamin 2. Data are means ± SEM (n ≥ 789 aggregates in 10 cells). (D) Average number of aggregates formed by either wild type or mutant dynamin 2 per 100 μm^2^ of the cell area. Data are means ± SEM (n ≥ 967 aggregates in ≥10 cells, N=3) (E) Quantification of the short TLS (0.5 ≤ 2 *μ*m) formed in the presence of wild type or novel mutant dynamin 2. Data are means ± SEM (n ≥ 242 TLS in ≥ 6 cells, N ≥ 3).

### Correlation of defective membrane tubulation and clinical phenotypes of patients

We hypothesized that the aberrant membrane remodeling activity by mutated dynamin 2 were implication of pathogenicity by each variant and clinical severity of CNM patients. Thus, we analyzed correlation between short TLS formation (Figs 1 and 2) and the clinical parameters of the patients with *DNM2* variants (ages of disease onset, biopsy and disease duration) (Supplementary Table 1 and 2). Among these parameters, ages of the disease onset and short TLS formation represented a linear correlation with high correlation coefficient (r = −0.74) (Fig. 3A). In contrast, the disease duration and the short TLS formation were not correlated (r = −0.1) (Figs. 3B). Thus, the defective membrane remodeling can explain and predict the variant-dependent occurrences of muscular weakness in *DNM2*-associated CNM. Importantly, P16 and P17 were distributed far from the line on the graph (Fig. 3A). Based on these data together with the pathological features, we concluded that novel variants, G495R, V520G and G624V, but not P294L and R724H, are likely to be pathogenic variants. Interestingly, the results of *in cellulo* experiments and ages of biopsy were also well correlated as plotted on an exponential curve (r = −0.97) except for those with R465W variant (Figs. 3C). As mentioned above, typical *DNM2*-associated CNM has milder and slowly progressive symptoms and favorable prognosis, but the onset is at infantile to adolescence. Our patients with reported variants were compatible to those features (Fig.3D). In contrast, the patients with novel variants were remarkably late-onset as a few previous reports about late-onset *DNM2*-CNM patients (Fig. 3D).

**Figure 3.**
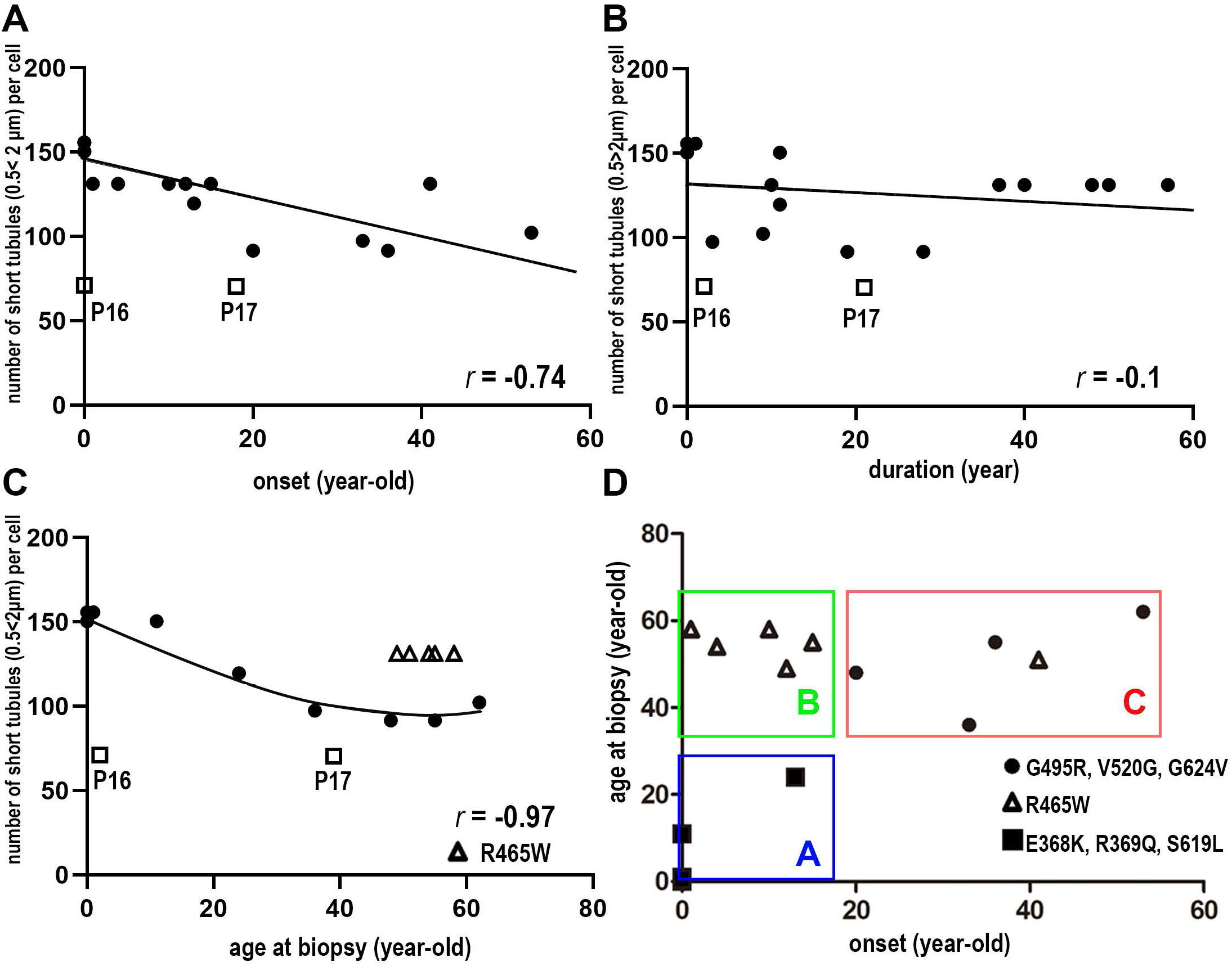
Correlation analyses between TLS formation and clinical phenotypes. Scatter plots for the presence of correlation between the short TLS formation and either disease onset age (A), disease duration (B) or age at biopsy (C). r indicates correlation coefficient. Non-pathogenic variants were shown with patient No in white square (P16 and P17). (D) Relationship between age at biopsy and onset of the disease. Group A, B and C represent severe phenotype (early onset and age at biopsy), slowly progressive phenotype (early onset and late biopsy) and late onset phenotype (late onset and age at biopsy), respectively.

## DISCUSSION

Recent advancement of the massive parallel sequencing technologies provides us with an enormous amount of genomic data from patients of various congenital diseases. Not surprisingly, pathogenicity of many of the identified variants is unknown, which clearly shows the necessity of simple and fast methods for assaying the functional defects caused by each variant.

In this study, we evaluated the pathogenicity of CNM-associated or -unassociated *DNM2* variants by imaging TLS formation in cultured cells and demonstrated that short TLS formation *in cellulo* and the severity of symptoms are correlated with high correlation coefficient (Fig. 3). This result suggests that the *in cellulo* assay in combination with the genetical and clinicopathological diagnosis are powerful approaches not only to determine pathogenicity of the genetic variants in *DNM2*, but also to predict the disease severity. Since the imaging-based *in cellulo* assay is easily accessible but provides with highly reproducible and quantitative results, it may applicable to elucidate pathomechanisms of triadopathies accompanied with disorganized T-tubules (Dowling, Lawlor, & Dirksen, 2014).

We analyzed the *DNM2* variants identified from CNM patients in our cohort (Supplementary Table.1 and 2). Both reported (E368K, R369Q, R465W and S619L) and novel (G495R, V520G and G624V) variants formed abnormal aggregates and short TLS (Figs. 1 and 2). These *in cellulo* phenotypes suggest that the novel variants, like reported *DNM2* variants, are responsible for gain-of-function features in self-assembly and GTPase activity, both of which are essential for membrane fission by dynamin 2. Furthermore, association between the deficits in TLS and clinical symptom of patients suggests that p.G495R, p.V520G and p.G624V are likely to be pathogenic (Fig. 3). Interestingly, G495R locates in the hinge between stalk (middle domain) and PH domain, whereas V520G and G624 are located in PH domain flanking stalk region based on the structure of dynamin 1 monomer (Supplementary Fig. 1). Previous structural studies on dynamin 1 and 3 demonstrated that PH domain is flipped back to interact with stalk and GTPase domain to form inhibitory “closed” state which is released upon binding to membrane by the PH domain (Faelber et al., 2011; Reubold et al., 2015). Thus, it is possible that these novel CNM variants affect structures of PH and stalk domains to impair the regulation of GTPase activity causing constitutively active GTPase state. Future structural studies of these novel dynamin 2 mutants will explain detailed mechanisms that cause their gain-of-function features linked to CNM pathogenesis.

In this study, we demonstrated correlation between the short TLS formation and ages of the patients when their biopsy samples were collected (Fig.3C), suggesting possible prediction of the *DNM2* variant-associated prognosis of the disease. Interestingly, p.R465W variant revealed unique features in both biological and clinical aspects: the GTPase activity and BIN1-susceptibility of this mutant were similar to those of wild type, and the patients have early age (infant to adolescence) at onset, and the relatively late age at biopsy as classified in Group B (Fig. 3D). Further studies are required to elucidate the precise pathogenesis by the R465W variant.

*DNM2*-associated CNM represents variable range of clinical phenotype with association between genetic variants and clinical severities (Bohm et al., 2012). However, most of the reported variants are associated with either early-onset severe phenotype (e.g., E368K, R369Q and S619L) or early-onset but with relatively mild phenotype (e.g., R465W). In contrast, only a few patients have been reported to develop late-onset disease. In support of this notion, all the patients with the reported variants in our cohort had either early-onset severe phenotype with early muscle biopsy (Group A) or early-onset but slowly progressive phenotype with late muscle biopsy (Group B) phenotypes. In contrast, all of our novel variants, i.e., p.G495R, p.V520G and p.G624V, were associated with the late-onset phenotype with late muscle biopsy (Group C) (Fig. 3D). Although the patients in Group C were almost asymptomatic until the third to fourth decades of their life, progression of muscle weakness after the onset was relatively rapid (Supplementary Table 2).

Several therapeutic applications targeting mutated dynamin 2 have been developed on animal studies (Buono et al., 2018; Trochet et al., 2018). Since a clinical trial using investigational antisense medicine DYN101 are ongoing for *DNM2*-associated CNM (NCT04033159) (https://www.clinicaltrials.gov/ct2/show/NCT04033159), establishing accurate diagnosis of CNM patients is crucial. Our approach using simple *in cellulo* assay together with genetical and clinicopathological analyses should contribute to precise diagnosis, especially when muscle biopsy samples are unavailable for any reasons. Furthermore, from the therapeutic point of view, early diagnosis by our simple assay also help to determine the management of the patients.

## Acknowledgements

The authors are thankful to P. Guicheney (UPMC), P. De Camili (Yale University) and H. McMahon (MRC-LMB) for reagents.

## REFERENCES

Agrawal, P. B., Pierson, C. R., Joshi, M., Liu, X., Ravenscroft, G., Moghadaszadeh, B.,… Beggs, A. H. (2014). SPEG interacts with myotubularin, and its deficiency causes centronuclear myopathy with dilated cardiomyopathy. Am J Hum Genet, 95(2), 218–226. doi:10.1016/j.ajhg.2014.07.004

Al-Qusairi, L., & Laporte, J. (2011). T-tubule biogenesis and triad formation in skeletal muscle and implication in human diseases. Skelet Muscle, 1(1), 26. doi:10.1186/2044-5040-1-26

Antonny, B., Burd, C., De Camilli, P., Chen, E., Daumke, O., Faelber, K.,… Schmid, S. (2016). Membrane fission by dynamin: what we know and what we need to know. EMBO J, 35(21), 2270–2284. doi:10.15252/embj.201694613

Biancalana, V., Romero, N. B., Thuestad, I. J., Ignatius, J., Kataja, J., Gardberg, M.,… Laporte, J. (2018). Some DNM2 mutations cause extremely severe congenital myopathy and phenocopy myotubular myopathy. Acta Neuropathol Commun, 6(1), 93. doi:10.1186/s40478-018-0593-2

Bohm, J., Biancalana, V., Dechene, E. T., Bitoun, M., Pierson, C. R., Schaefer, E.,… Laporte, J. (2012). Mutation spectrum in the large GTPase dynamin 2, and genotype-phenotype correlation in autosomal dominant centronuclear myopathy. Hum Mutat, 33(6), 949–959. doi:10.1002/humu.22067

Buono, S., Ross, J. A., Tasfaout, H., Levy, Y., Kretz, C., Tayefeh, L.,… Cowling, B. S. (2018). Reducing dynamin 2 (DNM2) rescues DNM2-related dominant centronuclear myopathy. Proc Natl Acad Sci U S A, 115(43), 11066–11071. doi:10.1073/pnas.1808170115

Casar-Borota, O., Jacobsson, J., Libelius, R., Oldfors, C. H., Malfatti, E., Romero, N. B., & Oldfors, A. (2015). A novel dynamin-2 gene mutation associated with a late-onset centronuclear myopathy with necklace fibres. Neuromuscul Disord, 25(4), 345–348. doi:10.1016/j.nmd.2015.01.001

Chin, Y. H., Lee, A., Kan, H. W., Laiman, J., Chuang, M. C., Hsieh, S. T., & Liu, Y. W. (2015). Dynamin-2 mutations associated with centronuclear myopathy are hypermorphic and lead to T-tubule fragmentation. Hum Mol Genet, 24(19), 5542–5554. doi:10.1093/hmg/ddv285

Cowling, B. S., Prokic, I., Tasfaout, H., Rabai, A., Humbert, F., Rinaldi, B.,… Laporte, J. (2017). Amphiphysin (BIN1) negatively regulates dynamin 2 for normal muscle maturation. J Clin Invest, 127(12), 4477–4487. doi:10.1172/JCI90542

Dowling, J. J., Lawlor, M. W., & Dirksen, R. T. (2014). Triadopathies: an emerging class of skeletal muscle diseases. Neurotherapeutics, 11(4), 773–785. doi:10.1007/s13311-014-0300-3

Faelber, K., Posor, Y., Gao, S., Held, M., Roske, Y., Schulze, D.,… Daumke, O. (2011). Crystal structure of nucleotide-free dynamin. Nature, 477(7366), 556–560. doi:10.1038/nature10369

Ferguson, S. M., & De Camilli, P. (2012). Dynamin, a membrane-remodelling GTPase. Nat Rev Mol Cell Biol, 13(2), 75–88. doi:10.1038/nrm3266

Fujise, K., Okubo, M., Abe, T., Yamada, H., Nishino, I., Noguchi, S.,… Takeda, T. (2020). Mutant BIN1-Dynamin 2 complexes dysregulate membrane remodeling in the pathogenesis of centronuclear myopathy. J Biol Chem. doi:10.1074/jbc.RA120.015184

Hohendahl, A., Roux, A., & Galli, V. (2016). Structural insights into the centronuclear myopathy-associated functions of BIN1 and dynamin 2. J Struct Biol, 196(1), 37–47. doi:10.1016/j.jsb.2016.06.015

Jungbluth, H., & Gautel, M. (2014). Pathogenic mechanisms in centronuclear myopathies. Front Aging Neurosci, 6, 339. doi:10.3389/fnagi.2014.00339

Jungbluth, H., Wallgren-Pettersson, C., & Laporte, J. (2008). Centronuclear (myotubular) myopathy. Orphanet J Rare Dis, 3, 26. doi:10.1186/1750-1172-3-26

Kenniston, J. A., & Lemmon, M. A. (2010). Dynamin GTPase regulation is altered by PH domain mutations found in centronuclear myopathy patients. EMBO J, 29(18), 3054–3067. doi:10.1038/emboj.2010.187

Majczenko, K., Davidson, A. E., Camelo-Piragua, S., Agrawal, P. B., Manfready, R. A., Li, X.,… Dowling, J. J. (2012). Dominant mutation of CCDC78 in a unique congenital myopathy with prominent internal nuclei and atypical cores. Am J Hum Genet, 91(2), 365–371. doi:10.1016/j.ajhg.2012.06.012

Marks, B., Stowell, M. H., Vallis, Y., Mills, I. G., Gibson, A., Hopkins, C. R., & McMahon, H. T. (2001). GTPase activity of dynamin and resulting conformation change are essential for endocytosis. Nature, 410(6825), 231–235. doi:10.1038/35065645

Nishikawa, A., Mitsuhashi, S., Miyata, N., & Nishino, I. (2017). Targeted massively parallel sequencing and histological assessment of skeletal muscles for the molecular diagnosis of inherited muscle disorders. J Med Genet, 54(2), 104–110. doi:10.1136/jmedgenet-2016-104073

Okubo, M., Iida, A., Hayashi, S., Mori-Yoshimura, M., Oya, Y., Watanabe, A.,… Nishino, I. (2018). Three novel recessive DYSF mutations identified in three patients with muscular dystrophy, limb-girdle, type 2B. J Neurol Sci, 395, 169–171. doi:10.1016/j.jns.2018.10.015

Ramachandran, R., Surka, M., Chappie, J. S., Fowler, D. M., Foss, T. R., Song, B. D., & Schmid, S. L. (2007). The dynamin middle domain is critical for tetramerization and higher-order self-assembly. EMBO J, 26(2), 559–566. doi:10.1038/sj.emboj.7601491

Reubold, T. F., Faelber, K., Plattner, N., Posor, Y., Ketel, K., Curth, U.,… Eschenburg, S. (2015). Crystal structure of the dynamin tetramer. Nature, 525(7569), 404–408. doi:10.1038/nature14880

Romero, N. B. (2010). Centronuclear myopathies: a widening concept. Neuromuscul Disord, 20(4), 223–228. doi:10.1016/j.nmd.2010.01.014

Schindelin, J., Arganda-Carreras, I., Frise, E., Kaynig, V., Longair, M., Pietzsch, T.,… Cardona, A. (2012). Fiji: an open-source platform for biological-image analysis. Nat Methods, 9(7), 676–682. doi:10.1038/nmeth.2019

Trochet, D., Prudhon, B., Beuvin, M., Peccate, C., Lorain, S., Julien, L.,… Bitoun, M. (2018). Allele-specific silencing therapy for Dynamin 2-related dominant centronuclear myopathy. EMBO Mol Med, 10(2), 239–253. doi:10.15252/emmm.201707988

Wang, L., Barylko, B., Byers, C., Ross, J. A., Jameson, D. M., & Albanesi, J. P. (2010). Dynamin 2 mutants linked to centronuclear myopathies form abnormally stable polymers. J Biol Chem, 285(30), 22753–22757. doi:10.1074/jbc.C110.130013

Warnock, D. E., Hinshaw, J. E., & Schmid, S. L. (1996). Dynamin self-assembly stimulates its GTPase activity. J Biol Chem, 271(37), 22310–22314. Retrieved from https://www.ncbi.nlm.nih.gov/pubmed/8798389

