## Supplementary Materials for "Imaging-based evaluation of pathogenicity by novel *DNM2 variants* associated with centronuclear myopathy"

#### *SUPPLEMENTARY METHODS*

##### *Genetic sequencing and histological analyses*

Genetic variants were analysed using genomic DNA from serum or biopsied muscles from 3933 cases which was suspected to be muscle diseases. Genetic analyses were performed by targeted re-sequencing that covered all exonic regions and exon-intron borders in *DNM2* gene (2858 analyses) and/or by whole exome sequencing (2599 analyses) as described previously [1]. Histological analysis was performed using muscle samples taken from the biceps brachii and then frozen in isopentane cooled in liquid nitrogen as described previously [2].

#### ***In vitro*** **sedimentation assay**

*In vitro* sedimentation assay of dynamin 2 was performed as described previously [3]. In short, wild type or CNM mutant (E368K, R369Q, R465W and S619L) dynamin 2 were diluted to 1 $\mu$M in reaction buffer (10 mM Hepes, 2 mM $MgCl_2$, 100 mM NaCl, pH 7.5) and incubated for 5 min at 37 °C. To induce disassembly, 1 mM GTP was added to the preassembled dynamin 2 and incubated for 5 min at 37 °C. The samples were centrifuged at 230,000*g* for 10 min at 25 °C using CS100GXL ultracentrifuge and S120AT3 rotor (Eppendorf Himac Technologies) and resultant supernatant and pellet were analyzed by

SDS-PAGE followed by Coomasie Brilliant Blue R-250 staining.

***Dynamin GTPase activity***

GTPase activity of dynamin 2 was determined by monitoring release of free orthophosphate using malachite green assay as described previously [3]. The malachite green reagent was prepared by mixing solution A (17 mg of Malachite Green Carbinol base dye (229105, Merck) in 20 mL 1 N HCl) and Solution B (0.5 g Ammonium molybdate (277908, Merck) in 7 mL 4 N HCl) with filling up to 50 mL by MilliQ water followed by filtration through 0.45 $\mu$m membrane (S-2504, KURABO). In the assay, 0.2 $\mu$M dynamin in the presence of BIN1 at different molar ratio was mixed with 1 mM GTP in GTPase reaction buffer (10 mM Hepes, 2 mM $MgCl_2$, 50 mM NaCl, pH 7.5) with or without 0.005 $\mu$g/$\mu$L lipid nanotubes and incubated for 5 min at 37 °C. After the reaction was stopped on ice for 10 min, 160 $\mu$L of malachite green reagent was added to the 40 $\mu$L of the reaction mix in 96 well plate (442404, Thermo Fisher Scientific). After 5 min shaking at 1200 rpm with Digital MicroPlate Genie Pulse (Scientific Industries, Inc.), released orthophosphate was colorimetrically quantified by measuring OD 650 nm using a microplate reader (SH-1000, CORONA ELECTRIC).

### Supplementary Figures

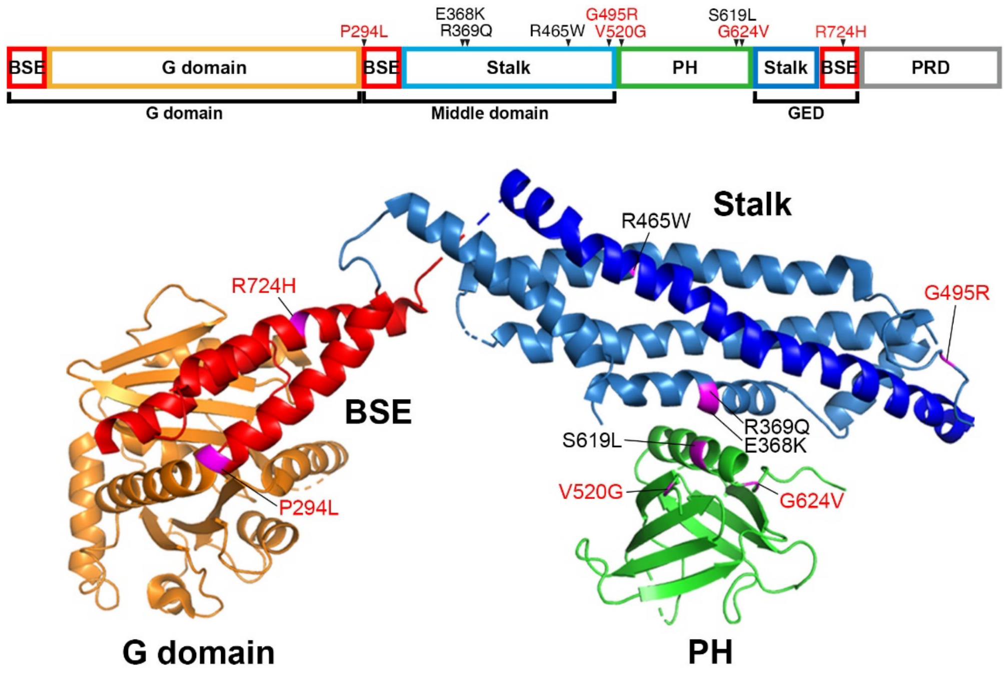


**Supplementary Figure 1 Location of mutations on dynamin.** (Upper) Schematic illustrations of dynamin 2 variants identified in this study. Previously reported and novel variants are shown in black and red, respectively. (Lower) Mutation sites are assigned on the crystal structure of human dynamin 1 (Protein Data Bank ID: 3SNH).

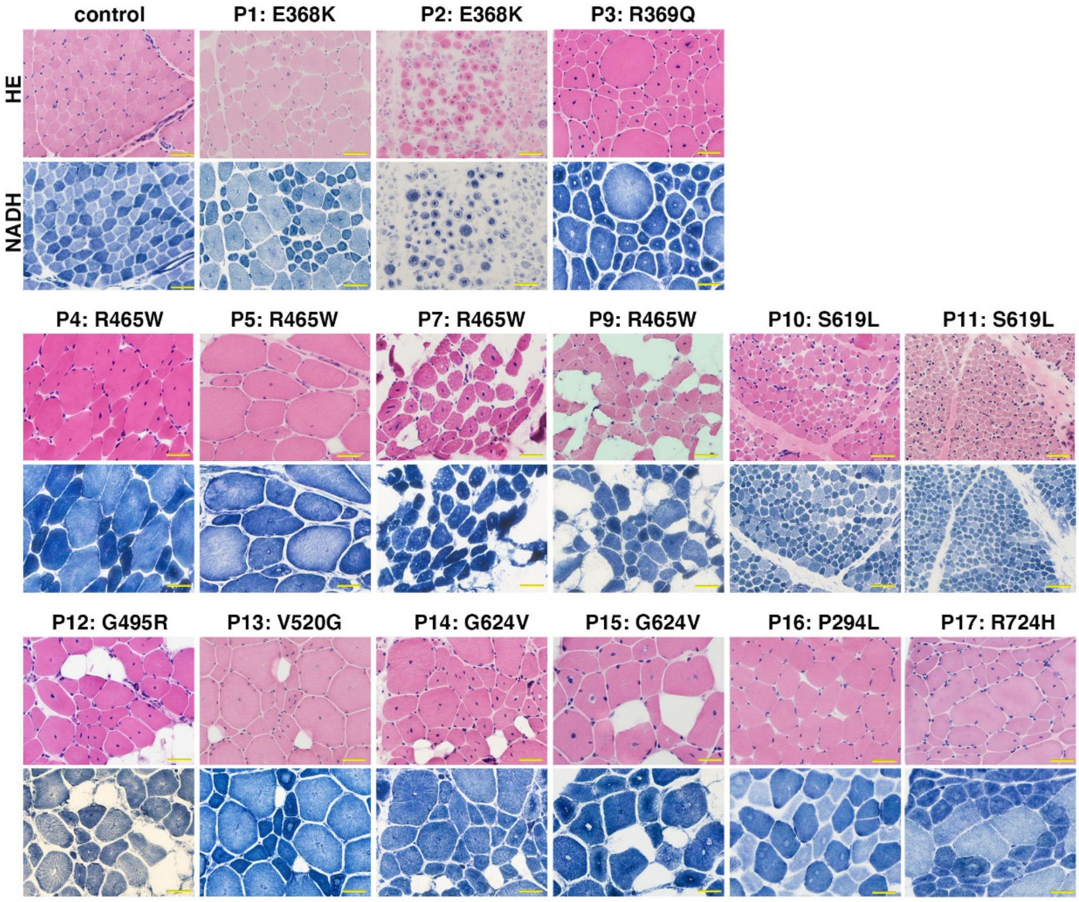


**Supplementary Figure 2 Histology of skeletal muscle biopsies from the patients with *DNM2* variants.** Staining with hematoxylin and eosin (HE, upper panels) and staining for nicotinamide adenine dinucleotide tetrazolium reductase (NADH, lower panels) are shown. The muscles from Patients 1-15 exhibit typical CNM pathology, centrally positioned nuclei, variation in myofiber size and adipose tissue replacement, while those from Patients 16 and 17 do not. Nondiagnostic disease-control (control) exhibits almost normal features. Scale bars are 10 $\mu$m.

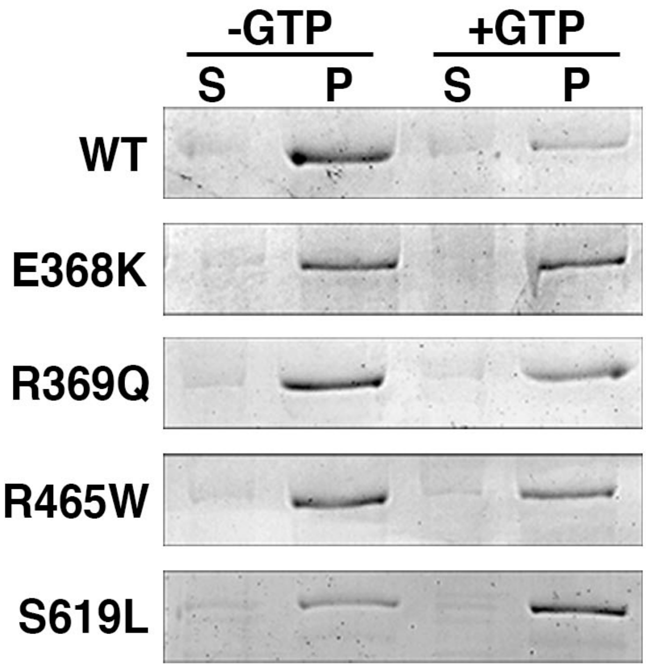


**Supplementary Figure 3 CNM mutant dynamin 2 form aggregates resistant to GTP hydrolysis.** CBB stained SDS-PAGE gel images for wild type (WT) or reported-mutant dynamin 2 (E368K, E369Q, R465W and S619L) fractionated in either the supernatant (S) or in the precipitate (P) with or without GTP (+GTP or -GTP) in the *in vitro* sedimentation assay.

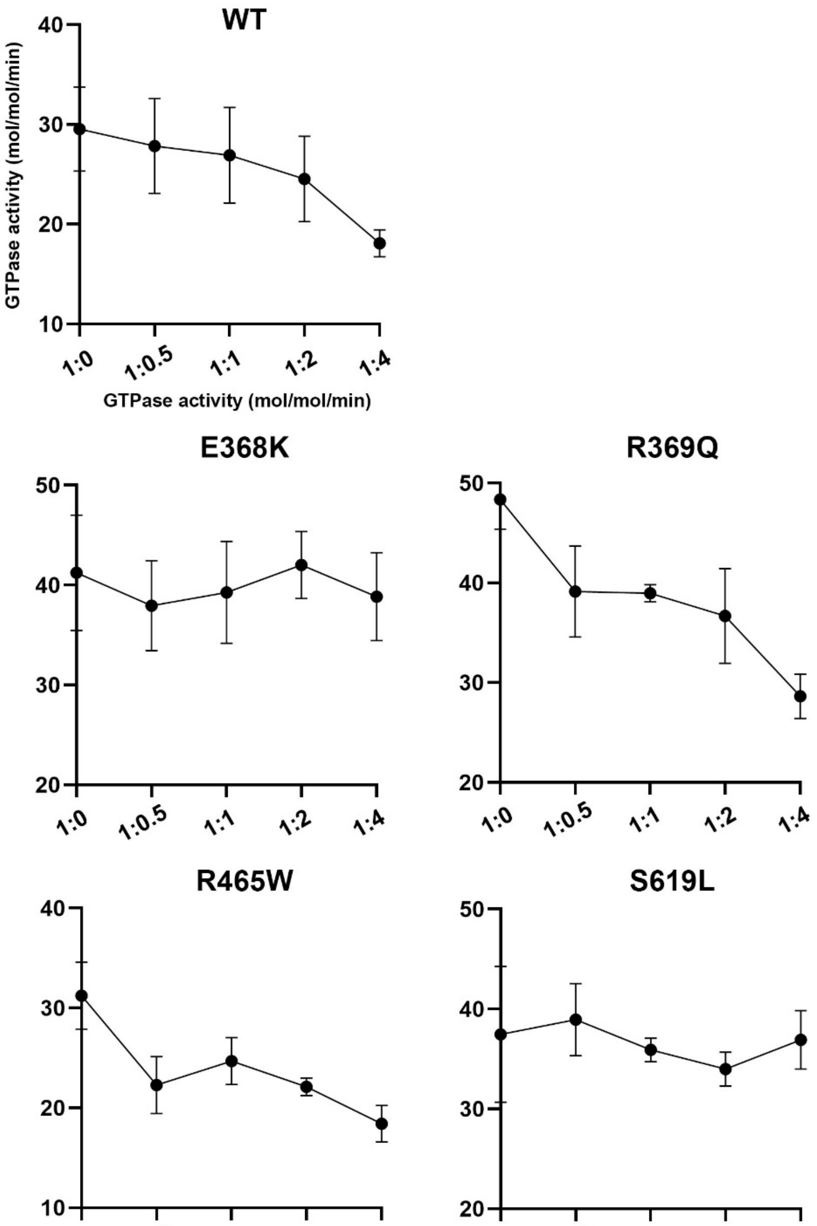


**Supplementary Figure 4 BIN1-mediaed suppression of GTPase activities of wild type and mutant dynamin 2.** Relative GTPase activities of either wild type (WT) and mutant dynamin 2 (E368K, R369Q, R465W and S619L) with increasing amount of BIN1 (1:0, 1:0.5, 1:1, 1:2 and 1:4 in molar ratio) are shown. Data are means ± SEM (n=3, N=3).

### *REFERENCES*


1. Nishikawa A, Mitsuhashi S, Miyata N, Nishino I. Targeted massively parallel sequencing and histological assessment of skeletal muscles for the molecular diagnosis of inherited muscle disorders. *J Med Genet*. Feb 2017;54(2):104-110. doi:10.1136/jmedgenet-2016-104073
2. Okubo M, Iida A, Hayashi S, et al. Three novel recessive DYSF mutations identified in three patients with muscular dystrophy, limb-girdle, type 2B. *J Neurol Sci*. Dec 15 2018;395:169-171. doi:10.1016/j.jns.2018.10.015
3. Fujise K, Okubo M, Abe T, et al. Mutant BIN1-Dynamin 2 complexes dysregulate membrane remodeling in the pathogenesis of centronuclear myopathy. *J Biol Chem*. Nov 13 2020;doi:10.1074/jbc.RA120.015184
