## Supplementary Tables for "Imaging-based evaluation of pathogenicity by novel *DNM2 variants* associated with centronuclear myopathy"

### Supplementary Table 1 Clinicopathological features of the patients with a reported mutation in *DNM2*

| **Patient** | P1 | P2 | P3 | P4 | P5 | P6 | P7 | P8 | P9 | P10 | P11 |
|---|---|---|---|---|---|---|---|---|---|---|---|
| **mutation** | c.1102G>A | c.1102G>A | c.1106G>A | c.1393C>T | c.1393C>T | c.1393C>T | c.1393C>T | c.1393C>T | c.1393C>T | c.1856C>T | c.1856C>T |
| **(AA change)** | (E368K) | (E368K) | (R369Q) | (R465W) | (R465W) | (R465W) | (R465W) | (R465W) | (R465W) | (S619L) | (S619L) |
| **Sex** | F | M | F | F | M | F | F | F | M | F | F |
| **age at last examination** | 11y | 0y2m | 24y | 55y | 51y | 49y | 58y | 58y | 54y | 1y5m | 0y5m |
| **Onset** | birth | birth | 13y | 15y | 41y | 12y | 1y | childhood | 4y | birth | birth |
| **Classification** | A | A | A | B | B | B | B | B | B | A | A |
| **Clinical presentation** | | | | | | | | | | | |
| Motor development | | | | | | | | | | | |
| head control | 5m | not acquired | normal | normal | normal | normal | normal | normal | normal | 1y | not acquired |
| age sitting | 6m | not acquired | normal | normal | normal | normal | normal | normal | normal | not acquired | not acquired |
| age ambulation | 4y | not acquired | 1y | 1y | 1y | 1y1m | 1y | normal | 1y2m | not acquired | not acquired |
| Ptosis | + | - | - | - | - | - | + | - | + | - | - |
| Limitation of eye movement | + | - | - | - | - | - | - | + | - | - | - |
| Facial weakness | - | + | - | + | - | + | - | - | - | + | + |
| High arched palate | - | + | + | + | - | - | - | - | - | + | + |
| Neck flexor weakness | ND | + | - | + | - | + | + | + | + | + | + |
| Respiratory disorder | - | + | + | + | + | - | + | + | + | - | - |
| Pattern of weakness | proximal | proximal | distal | diffuse | lower limb* | proximal | lower limb* | lower limb* | diffuse | diffuse | diffuse |
| age at not ambulatory | 23y | - | - | 50y | - | 48y | 50s | 47y (stick) | 49y | - | - |
| Scoliosis | - | - | - | - | - | - | - | - | - | - | - |
| Contractures | Achilles | knee | - | - | - | Achilles | - | Shoulder, Achilles | - | - | - |
| **Examination results** | | | | | | | | | | | |

| | | | | | | | | | | | |
|---|---|---|---|---|---|---|---|---|---|---|---|
| Serum CK (U/l) | 33 | 108 | 77 | 56 | 270 | 42 | 35 | 35 | 58 | 51 | 50 |
| **Muscle biopsy** | | | | | | | | | | | |
| central nuclei | + | + | + | + | + | + | + | ND | + | + | + |
| type 1 fiber predominance | - | - | + | + | + | + | + | ND | - | + | - |
| Radial strands | + | + | + | + | + | + | + | ND | + | - | + |

*muscle weakness was observed in lower limb, but the pattern of weakness was not described.

### Supplementary Table 2 Clinicopathological features of the patients with a novel variant in *DNM2*

| **Patient** | P12 | P13 | P14 | P15 | P16 | P17 |
|---|---|---|---|---|---|---|
| **Variant (AA change)** | c.1483G>A (G495R) | c.1559T>G (V520G) | c.1871G>T (G624V) | c. 1871G>T (G624V) | c.881C>T (P294L) | c.2171G>A (R724H) |
| **Sex** | M | F | M | M | F | F |
| **age at last examination** | 62y | 36y | 48y | 55y | 39y | 2y |
| **Onset** | 53y | 33y | 20y | 36y | 18y | birth |
| **Classification** | C | C | C | C | NP** | NP** |
| **Clinical presentation** | | | | | | |
| Motor development | | | | | | |
| head control | normal | normal | normal | normal | normal | 6m |
| age sitting | normal | normal | normal | normal | normal | not acquired |
| age ambulation | normal | normal | normal | normal | 1y | not acquired |
| Ptosis | - | - | - | - | - | - |
| Limitation of eye movement | - | - | - | - | - | - |
| Facial weakness | - | - | ND | - | - | + |
| High arched palate | - | - | ND | - | - | + |
| Neck flexor weakness | ND | + | ND | - | - | + |
| Respiratory disorder | + | ND | - | - | - | - |
| Pattern of weakness | proximal | lower limb* | distal | distal | lower limb proximal | proximal |
| age at not ambulatory | - | - | - | - | - | - |
| Scoliosis | - | - | - | - | - | - |
| Contractures | - | - | - | - | - | - |
| **Examination results** | | | | | | |

| Serum CK (U/l) | 223 | 116 | 97 | 87 | 68 | 60 |
|---|---|---|---|---|---|---|
| **Muscle biopsy** | | | | | | |
| central nuclei | + | + | + | + | - | + |
| type 1 fiber predominance | - | - | + | - | - | + |
| Radial strands | + | + | + | + | - | - |

*muscle weakness was observed in lower limb, but the pattern of weakness was not described.

**NP: non-pathogenic variant
